# Partial cross-resistance of oats to different *Fusarium* species and the role of trichomes in susceptibility

**DOI:** 10.1101/2025.04.08.647721

**Authors:** Selma Schurack, Charlotte Rodemann, Klaus Oldach, Steffen Beuch, Sophie Brodführer, Andreas von Tiedemann, Matthias Herrmann

**Affiliations:** Institute for Breeding Research on Agricultural Crops, Julius Kühn Institute (JKI), Federal Research Centre for Cultivated Plants, Sanitz, Germany; Plant Pathology and Crop Protection Division, Georg-August University Goettingen, Goettingen, Germany; KWS LOCHOW GmbH, Bergen, Germany; Nordsaat Saatzucht GmbH, Zuchtstation Granskevitz, Schaprode, Germany; Saatzucht Bauer GmbH & Co. KG, Obertraubling, Germany

**Author notes:** Correspondence: **Matthias Herrmann**.

**Keywords:** Fusarium head blight, oats, Fusarium graminearum, Fusarium poae, Fusarium sporotrichioides, qPCR, resistance, trichomes

## Abstract

Resistance of oats to FHB, caused by different *Fusarium* species, is important for grain quality and yield. In this study, 25 oat genotypes were evaluated for resistance to *Fusarium graminearum* (FG), *F. sporotrichioides* (FS) and *F. poae* (FP) in field trials across Germany to assess the presence of cross-resistance and to analyse the role of trichomes on the hulls during *Fusarium* infection. Infection severity was quantified by *Fusarium* species-specific qPCR and showed the highest fungal biomass for FP, followed by FS and FG. Variability due to environmental effects was very high, resulting in rather low heritabilities for FG (0.50) and FS (0.36), and no significant genotype effect for FP. A significant positive correlation was found between FP and FS infection, whereas FG infection was not correlated with either FP or FS. Microscopic analyses revealed important genotype-specific differences in trichome size and density on lemma and palea with very high heritability (0.97). FG biomass was positively correlated with trichome size and density, and FG hyphae were observed in close interaction with trichomes and stomata. These results indicate the presence of partial cross-resistance in addition to mostly species-specific resistance and suggest a role for trichomes in susceptibility to FG.

## 1 Introduction

Oats (*Avena sativa* L.) are well known for their beneficial nutritional properties and positive effects on soil health, but like other small grain cereals such as wheat and barley, they are susceptible to *Fusarium* head blight disease (FHB) caused by several *Fusarium* species (Placinta et al., 1999; Bottalico and Perrone, 2002). FHB in oats can lead to yield losses, reduced seed germination and the accumulation of mycotoxins such as deoxynivalenol (DON), nivalenol (NIV), T-2 and HT-2 which are harmful to human and animal health (Scott, 1989; Bjørnstad and Skinnes, 2008; Tekle et al., 2013). The spectrum of *Fusarium* species on oats and the severity of infection are subject to strong regional and annual variation, depending on the previous crop, tillage practices, the proportion of cereals and maize in the rotation and weather conditions (Bottalico et al., 2002; Hofgaard et al., 2016a; Parikka et al., 2012).

A recent survey of the occurrence of *Fusarium* species in oats in Germany identified the presence of *F. poae*, *F. tricinctum*, *F. avenaceum*, *F. culmorum*, *F. equiseti*, *F. graminearum*, *F. sporotrichioides*, *F. langsethiae* and *F. cerealis*. The most prevalent species was *F. poae*, followed by *F. tricinctum* and *F. avenaceum* (Rodemann et al., 2023). Similar species occurrences and a dominance of *F. poae*, *F. graminearum*, *F. avenaceum* and *F. langsethiae* were observed in other studies conducted across Central Europe (Nielsen et al., 2011; Kiecana et al., 2012; Dal Prá et al., 2014; Vanheule et al., 2014; Georgieva et al., 2018; Schöneberg et al., 2018). In Scandinavia, Finland and Canada, *F. avenaceum*, *F. culmorum* and *F. sporotrichioides* were also very common (Fredlund et al., 2013; Hietaniemi et al., 2016; Hofgaard et al., 2016b; Tekauz et al., 2008).

The co-occurrence of several *Fusarium* species raises the question of whether it is necessary to breed for resistance to each important species individually, or, ideally, whether generally effective, species non-specific FHB resistances are present in oats (i.e. cross-resistance). In wheat and barley, several studies have shown the broad effectiveness of different sources of resistance for different *Fusarium* species (Akinsanmi et al., 2006; Holzknecht et al., 2009; Mesterházy et al., 1999, 2005; Šíp et al., 2011; Tóth et al., 2008; van Eeuwijk et al., 1995). Nevertheless, there is still a lack of knowledge about the numerous other toxins that occur in addition to DON and that are relevant for consumer and animal protection (Mesterhazy, 2024). Further QTL analyses with segregating populations and inoculation with additional *Fusarium* species are required to validate the species-non-specific resistance observed in wheat (Mesterhazy, 2024). In oats, the situation regarding the species-specificity of resistance to FHB is less well understood. Some studies indicate that resistance to FHB is rather species-specific in oats (Aamot et al., 2022; Hofgaard et al., 2022; Tekauz et al., 2004). The clearest example is the variety Odal, which is relatively resistant to DON but is very susceptible to T-2/HT-2 accumulation (Hofgaard et al., 2022). However, Herrmann et al. (2020) found a modest but significant correlation between the ranking for T-2/HT-2 and DON contamination and a rather weak correlation between DON and T-2 levels in another unpublished series of experiments on modern variety material. In consideration of this, the existence of some species-unspecific resistance (i.e. cross-resistance) in addition to species-specific resistance cannot be ruled out. Further research is therefore needed to clarify this issue.

Resistance of small grain cereals against FHB is a complex phenomenon with several passive and active components (Mesterházy, 1995). Active defence responses include reinforcement of cell walls, counteracting oxidative stress or detoxifying mycotoxins, which can be virulence factors in wheat or barley (Lemmens et al., 2005; Spanic et al., 2017; Kheiri et al., 2019; Soni et al., 2021; Khairullina et al., 2022; Bethke et al., 2023). A few passive, morphology-based FHB resistance components have been identified in oats so far, such as plant height, epicuticular wax layer or anther extrusion (He et al., 2013; Loskutov et al., 2017; Hautsalo et al., 2020; Herrmann et al., 2020; Tekle et al., 2020; Willforss et al., 2020). As the primary interface for FHB infection is usually the outer parts of the flower where the *Fusarium* spores germinate, anatomical features of the hulls, such as trichomes, could also play an important role in infection in oats (Tekle et al., 2012). Trichomes are hair-like structures found on the aerial organs of most terrestrial plants and play different roles in plant development and stress response (Zhang et al., 2021; Han et al., 2022). It has been demonstrated that trichomes represent a preferred entry site for *Fusarium* hyphae in barley, maize, wheat and *Brachypodium* (Imboden et al., 2018; Linkmeyer et al., 2013; Peraldi et al., 2011; Sun et al., 2024). Similarly, in several wild *Avena* species, trichome abundance was positively correlated with DON and *Fusarium* biomass (Gagkaeva et al., 2017). However, also the opposite, where a higher trichome density was associated with FHB resistance, has been observed (Duba et al., 2019). In cultivated oats, the genetic variability of trichome abundance on the hulls and its relationship to FHB resistance has yet to be investigated.

The objectives of this study were (I) to assess resistance to three *Fusarium* species (*F. graminearum, F. sporotrichioides* and *F. poae*) to gain further insight into species-specific and cross-resistance mechanisms and (II) to characterize trichome variations on the surface of the hulls and to investigate their role during *Fusarium* infection in a panel of mostly modern German oat varieties.

## 2 Material and methods

### 2.1 Plant material

The studied oat panel consisted of 25 spring oat genotypes with similar plant heights and panicle emergence dates to limit the effect of these traits on potential differences in *Fusarium* resistance. Most genotypes were modern German varieties. Supplementary Table S1 provides a detailed description of the panel.

### 2.2 Fungal isolates and inoculum production

The fungal isolates used in this study comprised four *F. poae*, four *F. graminearum* and two *F. sporotrichioides* isolates derived from field samples of oats and maize from different sites in Germany and Austria. We selected these three *Fusarium* species because they were found to be highly prevalent in German oat fields in a previous monitoring project (Georgieva et al., 2018) and because they belong to the main producers of the three important mycotoxins DON (*F. graminearum*), T-2/HT-2 (*F. sporotrichioides*) and NIV (*F. poae*). A detailed description of the fungal isolates is given in Supplementary Table S2. Fungal isolates were grown on potato dextrose agar (PDA) plates at room temperature for approximately 7 days. SacO_2_ Autoclave Bags (MycoGenetics, Münster, Germany) filled with approximately 1.1 kg of oat kernels and 1 L ddH_2_O were autoclaved twice. After cooling, the bags were inoculated with small pieces of PDA containing mycelium of each isolate for each *Fusarium* species separately. The bags were incubated at room temperature for 1-2 weeks until all kernels were colonised. Then, the colonised kernels were spread in trays to allow further fungal development and to dry for 2-3 weeks. After that, the inoculum was collected and stored at – 20 °C until inoculation of the field trials. For spray-inoculation, liquid inoculum was prepared from the colonized kernels as described in Linkmeyer et al. (2013). In brief, colonised kernels were incubated in 0.02 % Tween20 in tap water for 15 min. The conidia suspension was filtered through filter paper, sedimented overnight and the supernatant was stored at 4 °C until usage.

### 2.3 Field trials

The oat panel was grown in three locations across Germany (Groß Lüsewitz (GL; coordinates <u>54.0714, 12.3238</u>), Wohlde (WL; coordinates 52.8074, 10.0003), Böhnshausen (BH, coordinates 51.8596, 10.9611) in the years 2021 and 2022. The genotypes were sown in plots of 1.8 sqm with a density of 350 kernels/sqm in a randomised complete block design with three replications for each *Fusarium* species. The trials for the different *Fusarium* species were planted with at least 9 m distance to avoid cross-contamination. Each plot was inoculated with 50 ml colonised kernels at BBCH 35±5 in 2021 (May 28 to June 8) and BBCH 55±5 in 2022 (June 13 and 14) (Meier, 2018). Plots were harvested either by a combine harvester or by hand, cleaned after threshing using the same wind sifter and after cleaning, seeds were analysed for *Fusarium* biomass.

### 2.4 DNA extraction and quantification of fungal biomass

Fungal DNA of *F. graminearum*, *F. sporotrichioides* and *F. poae* was quantified using quantitative real-time PCR (qPCR). Kernels were lyophilized and finely ground to a size of approx. 1 mm using a swing mill (Retsch MM 400, Retsch, Haan, Germany). DNA extraction was performed by using a CTAB-based extraction protocol described previously by Guerra et al. (2020). Quality and quantity of DNA extracts were assessed on agarose gels (0.8% (w/v) in 1x TAE (Tris-acetate-EDTA buffer) stained with Midori Green (Nippon Genetics Europe, Düren, Germany). Before qPCR analysis samples were diluted 1:50 in double-distilled H_2_O (ddH_2_O). DNA standards for qPCR assays were obtained from *F. poae* DSM62376, *F. graminearum* IFA66 und *F. sporotrichioides* DSM62423.

DNA for quantification standards were extracted using a CTAB-based protocol (Brandfass and Karlovsky, 2008) and quantified by densitometry (Nutz et al., 2011). PCR analysis was carried out using a CFX 384 Thermocycler (Biorad, Rüdigheim, Germany) in 384-well microplates. The reaction was carried out with 1 µL diluted DNA extracts in 4 µL reaction volume. Previously published species-specific primers were used (Supplementary Table S3, Nicholson et al., 1998; Parry et al., 1996; Wilson et al., 2004). All final concentrations of the used reagents are given in Supplementary Table S4. All standards and negative controls (ddH_2_O) were amplified in duplicates. Standard curves were generated from 1:3 serial dilutions of 100 pg/µL fungal DNA. PCR conditions and quantification limits (LOQ) are given in Supplementary Table S5. After amplification, melting curves were generated by increasing temperature from 55 °C to 95 °C with 0.5 °C increase per step and continuous fluorescence measurement. Samples with concentrations of fungal DNA below the LOQ but with positive melting curves were set to half of the LOQ (Clarke, 1998).

### 2.5 Analysis of trichome variation

To assess variation in hull trichome size and density, the lemma and palea of the 25 oat genotypes studied for *Fusarium* cross-resistance were examined. For this, samples from the field in Groß Lüsewitz in 2021 and 2022, as well as samples from plants grown in the greenhouse in 2021 and 2022 were used (= four environments). The lemma and palea from six individual plants per environment were collected at anthesis and fixed in 96 % EtOH. EtOH was replaced by fresh EtOH several times until all chlorophyll had been removed from the tissue. Three images per hull were captured using a Nikon Eclipse 90i fluorescence microscope with the standard series Nikon DAPI filter block (Excitation 375/28 nm, Barrier 460/60 nm). The cell counter plug-in in ImageJ (Vos, 2001; Schneider et al., 2012) was used to count the number of small (< 25 µm), medium (25 – 100 µm) and big (> 100 µm) trichomes on the exterior of the hulls (see Supplementary Figure S2 for example picture). A trichome index was calculated using the following formula: *Index = n. ts + 2 ∗ n. tm + 4 ∗ n. tb* with n.ts: number of small trichomes, n.tm: number of medium trichomes and n.tb: number of big trichomes. Weighing factors were chosen to take trichome size into account.

### 2.6 Confocal laser-scanning microscopy

To visualise fungal growth on the lemma and palea, oat plants grown in the greenhouse were spray-inoculated with *F. graminearum* (5×10^4^ conidia per ml) at anthesis. The oat varieties used were ‘Gabriel’, ‘Contender’ and ‘Max’. Inoculated panicles were covered with clear plastic bags for 48 h. Samples were collected at 24 hpi, 48 hpi and 96 hpi and fixed in 96 % EtOH. EtOH was replaced by fresh EtOH several times until all chlorophyll had been removed from the tissue. The samples were stained with Wheat Germ Agglutinin, AF488 Labelled (WGA-AF488, AAT Bioquest, CA, USA), trypan blue (TB, Merck) and aniline blue diammonium salt (AB, Sigma-Aldrich Chemie, Traufkirchen, Germany) as described by Becker et al. (2018) with the following modifications: Before staining, lemma samples were incubated for 3 h and palea samples were incubated for 1 h in 1 M KOH at room temperature. Staining solution contained 0.02 % AB, 0.02 % TB, 0.0005 % WGA-AF488 and 0.02 % Tween20 in PBS (pH 7.4). Confocal laser scanning microscopy (CLSM) was done using a Leica TCS SP8 as described by Becker et al. (2018) with the following changes: AB, TB and WGA-AF488 were excited simultaneously with the 405 nm diode, 488 nm argon ion and 561 nm diode-pumped solid-state (DPSS) lasers. Four detectors were used in sequential acquisition mode to capture emission fluorescence from the dyes and plant autofluorescence: HyD1 (380 nm – 414 nm, grey pseudocolor, plant autofluorescence), PMT2 (447 nm – 481 nm, blue pseudocolor, emission from WGA-AF488), HyD3 (593 nm – 626 nm, green pseudocolor, plant autofluorescence and emission from AB) and PMT4 (674 nm – 749 nm, red pseudocolor, plant autofluorescence and emission from AB and TB).

### 2.7 Data analysis

All statistical data analyses were performed in R (version 4.1.0; R Core Team, 2021). *Fusarium* DNA contents were square root transformed for statistical analyses. BLUEs were estimated by fitting a mixed linear model with the ‘lmer’ function of the ‘lme4’ package (version 1.1.35.1; Bates et al., 2015). Genotype effects were set as fixed and environment, environment-genotype interaction and replication effects nested in the environment were set as random. For estimation of variance components, the same model was applied, but with setting genotype effects as random as well.

Significance of genotype effects was calculated using the ‘anova’ function (version 4.1.0; R Core Team, 2021). Broad-sense heritability was estimated with the formula according to Becker (2019). Spearman-rank correlation coefficients and respective p values were calculated using the ‘rcorr’ function in the ‘Hmisc’ package (version 4.7.2; Frank E Harrell Jr, 2022).

## 3 Results

### 3.1 Oat resistance to different *Fusarium* species

To assess resistance levels against different *Fusarium* species, field trials using 25 oat genotypes were separately inoculated with *F. graminearum* (FG), *F. sporotrichioides* (FS) and *F. poae* (FP) field isolate mixtures via spawn-inoculation. The trials were performed with three replicates per oat genotype and *Fusarium* species in three locations across Germany (BH: Böhnshausen, GL: Groß Lüsewitz, WL: Wohlde) in two years (2021, 2022). To minimize the effect of plant height and developmental stage on *Fusarium* infection, we used oat genotypes that were found to be highly similar in these traits in previous experiments. To verify this, plant height and panicle emergence date was recorded in Groß Lüsewitz in 2021. For plant height, means of the genotypes ranged from 78 to 88 cm, and panicle emergence occurred within a period of 3 days (Supplementary Table S7). Hence, as expected, variance in these traits was small in the studied oat panel.

Infection severity was assessed via quantification of fungal DNA in the harvested oat seeds by species-specific qPCR assays. This showed very high variations across replicates, locations and years and the observed values ranged from zero to 388 µg fungal DNA per kg of harvested seeds (Figure 1A; Table 1). In 2022, infection levels were extremely low across all locations, and only for FP notable DNA quantities were detected. This might be due to unfavourable weather conditions prevalent during infection in 2022. Hence, for further analyses, we only used the data from 2021. Across locations and genotypes, fungal DNA content was highest for FP (29.49 µg/kg), followed by FS (21.17 µg/kg), and lowest for FG (11.58 µg/kg; Table 1). For all three fungal species, the environment explained most of the observed variance. Accordingly, heritability was rather low for FS and FG (0.36 and 0.50 respectively, Table 1). For FP there was no significant genotype effect, hence heritability was zero.

**Figure 1.**
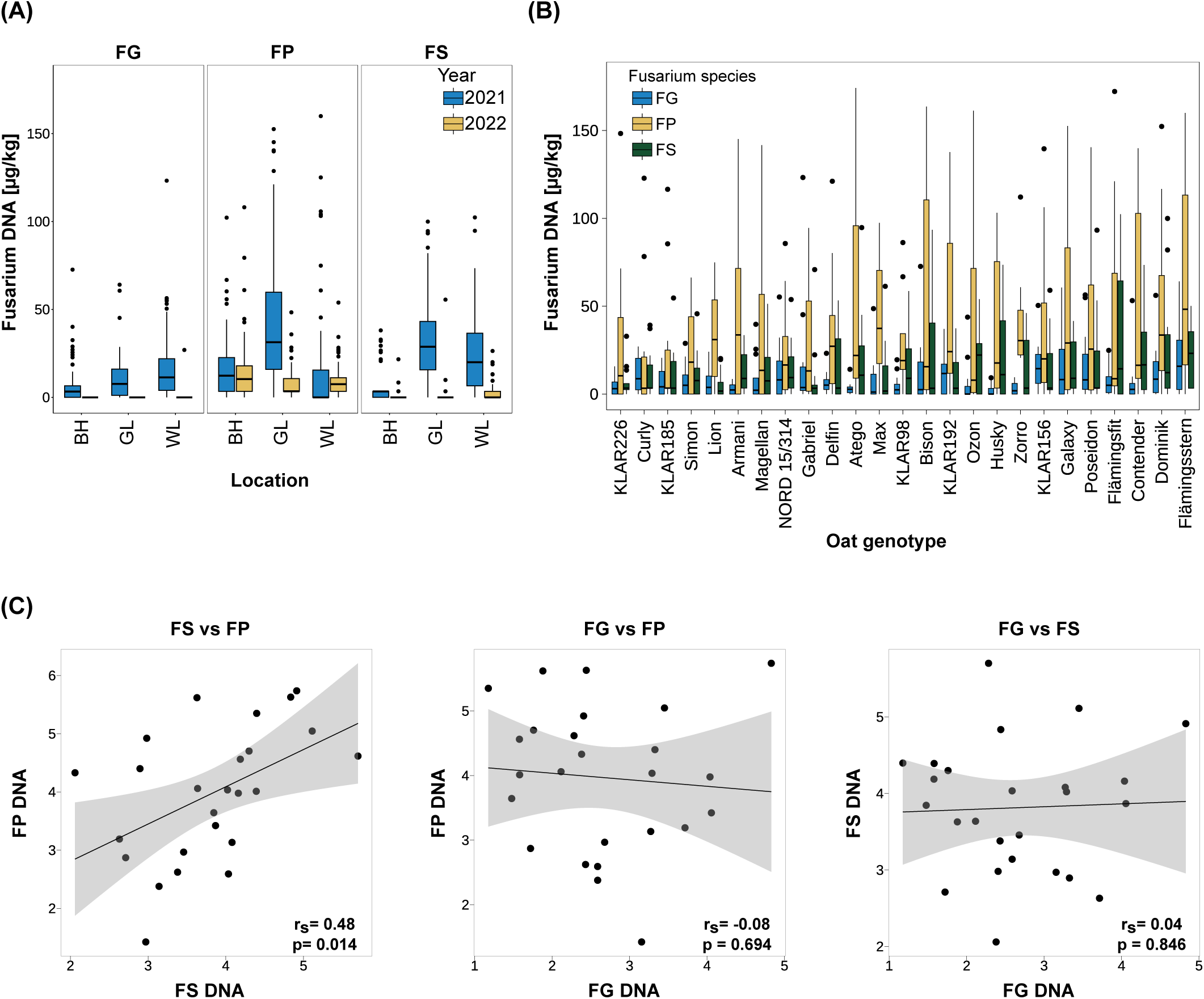
(A) *Fusarium* biomass in oat seeds across locations and years for different *Fusarium* species. Fungal biomass was assessed via quantification of fungal DNA per kg oat seeds. n=3 replicate plots per *Fusarium* species, oat genotype, study location and year. BH: Böhnshausen; GL: Groß Lüsewitz; WL: Wohlde. For visualization purposes, the y-axis limit was set to 180 µg/kg. (**B) *Fusarium* biomass across 25 oat genotypes for different *Fusarium* species.** Genotypes are ordered from low (left) to high (right) sum of fungal DNA of all three *Fusarium* species. Only 2021 data used. For visualization purposes, the y-axis limit was set to 180 µg/kg. (**C) Correlation of biomass of the different *Fusarium* species across oat genotypes.** BLUEs were used for calculation of Spearman rank correlation coefficients. Values were square root transformed before BLUE calculation. rs: Spearman-rank correlation coefficient; p: p-value. FG: *F. graminearum*; FP: *F. poae*; FS: *F. sporotrichioides*.

**Table 1.**
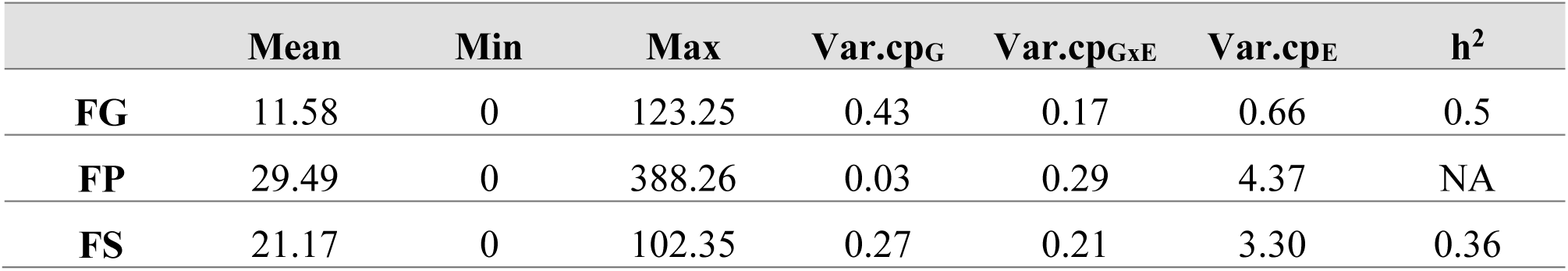
Means and variance components of *Fusarium* DNA content. Mean, minimum and maximum values are given for the untransformed data (given in µg fungal DNA per kg oat seeds), variance components are calculated based on the square root-transformed values. FG: *F. graminearum*; FP: *F. poae*; FS: *F. sporotrichioides*.

Across genotypes, mean FP infection was by far the highest in GL 2021 (56.04 µg/kg vs. 17.34 µg/kg in BH and 15.10 µg/kg in WL; Figure 1A, Supplementary Table S6). For FS, highest mean infection was as well observed in GL (33.3 µg/kg), while lowest mean infection was observed in BH (8.9 µg/kg; Figure 1A). FG mean infection was the highest in WL (17.47 µg/kg), followed by GL (10.62 µg/kg) and BH (6.64 µg/kg; Figure 1A, Supplementary Table S6).

The fungal DNA content in the harvested seeds of the tested oat genotypes for the different *Fusarium* species is shown in Figure 1B. The genotypes are ordered from lowest (left) to highest (right) total fungal DNA (sum of FG, FP and FS DNA of each genotype). In line with the overall highest mean, in most genotypes, DNA contents were highest for FP (12 genotypes), closely followed by FS (10 genotypes). Accordingly, in the majority of oat genotypes, FG DNA contents were the lowest compared to FS and FP (17 genotypes, Figure 1B).

To clarify whether species-specific resistance or species-unspecific resistance (i.e. cross-resistance) mechanisms were present, Spearman rank correlations between the different *Fusarium* species were calculated (Figure 1C). BLUEs were used for calculation of Spearman rank correlation coefficients and values were square root transformed before BLUE estimation. This identified a significant positive correlation of 0.48 between FP and FS infection severity. No significant correlations were detected between FG and FS or FP.

Despite the low correlation between the three *Fusarium* species, oat genotypes with good resistances to all three *Fusarium* species could be identified in the analysed oat panel (see genotypes on the left of Figure 1B, e.g. KLAR226, Curly, KLAR185, and NORD 15/314).

### 3.2 Variation of trichome size and density on oat hulls

To evaluate variations in surface morphology that could affect oat – *Fusarium* interactions, we examined differences in trichome size and density on the paleas and lemmas of the 25 oat genotypes in the cross-resistance panel. For this, lemma and palea samples were taken at anthesis from plants grown in the field and in the greenhouse in 2021 and 2022 (= four environments). Trichomes were visualized by autofluorescence using a fluorescence microscope and counted classified by size. All trichomes observed were of the prickle type, no macro hairs were noted. Trichome density generally decreased from tip to base of both organs. To summarize trichome size and density, a trichome index considering both values was calculated. Heritability of the trichome index was very high (0.97) and trichome density and size profiles differed strongly between the analysed oat genotypes (Figure 2A). Figure 2B shows exemplary paleas of the extreme genotypes Atego (lowest trichome index) and KLAR156 (highest trichome index). In general, trichome density and index was higher on the lemma than the palea, and most trichomes fell into the ‘medium’ category. Furthermore, trichome indices as well as trichome densities of lemma and palea showed a strong significant positive correlation (Spearman rank correlation coefficient 0.83 and 0.99, respectively, Supplementary Figure S1).

**Figure 2.**
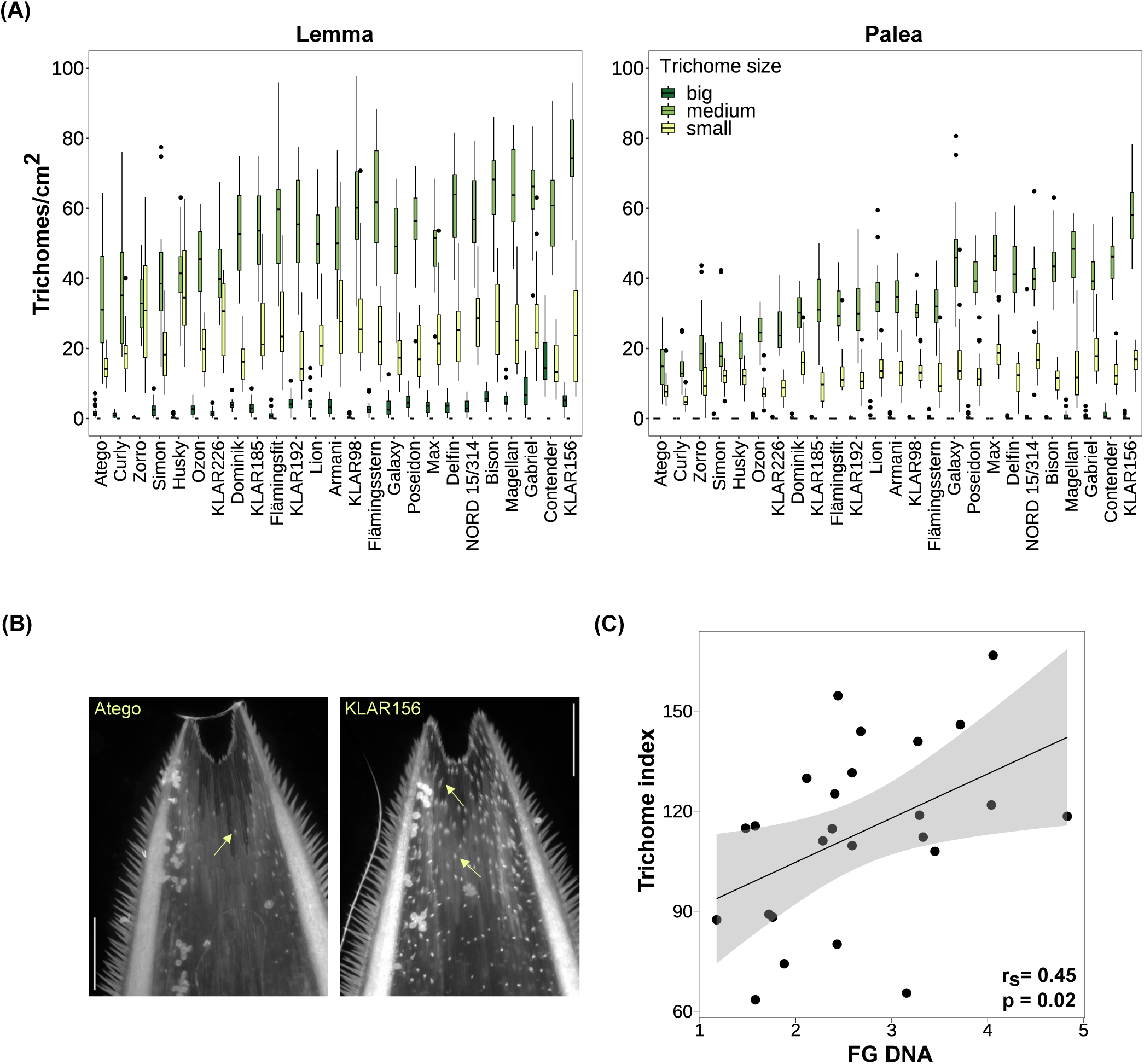
(A) Trichome density on lemma and palea across oat genotypes. Genotypes are ordered by trichome index from low (left) to high (right). Small trichomes: < 25 µm; medium trichomes: 25 – 100 µm; big trichomes > 100 µm. **(B) Exemplary paleas of oat genotypes with lowest (Atego) and highest (KLAR156) trichome index.** Trichomes were visualized through autofluores cence using the DAPI filter on a fluorescence microscope. Arrows indicate trichomes. Scale bar: 500 µm. **(C) Correlation of lemma and palea trichome index and *F. graminearum* biomass across oat genotypes.** BLUEs were used for calculation of Spearman rank correlation coefficients. FG: *F. graminearum*; rs: Spearman-rank correlation coefficient; p: p-value.

### 3.3 Relationship of trichome size and density and *Fusarium* infection

To understand whether the observed differences in trichome density and size profiles could play a role during *Fusarium* infection, we calculated their correlation with *Fusarium* biomass. For FG, a significant positive correlation with both trichome index and trichome density was found (r_s_=0.45 and 0.43, respectively; Figure 2C, Supplementary Figure S1). Furthermore, the individual values of trichome index and density of palea and lemma were also significantly positively correlated with FG biomass and showed similar correlation coefficients (r_s_= 0. 43 and 0.45, respectively, Supplementary Figure S1). The strongest correlation with FG biomass was observed with the density of medium-sized trichomes on lemmas (r_s_=0.53, Supplementary Figure S1). Furthermore, the trichome density of the five oat genotypes most susceptible to FG was about 1.6 times higher than that of the five oat genotypes most resistant to FG. In contrast, there was no significant correlation of any of the trichome measures with FS and FP (Supplementary Figure S1).

To gain further insight into the possible role of trichomes during infection, the interaction of FG hyphae with trichomes on the surface of paleas was investigated using confocal laser-scanning microscopy (Figure 3). Florets of the oat varieties ‘Gabriel’, ‘Contender’ and ‘Max’ were spray inoculated with FG conidia at anthesis and samples were taken at 24, 48 and 96 hpi. In several cases, hyphae grew in close contact with trichomes, often with accumulation of hyphae at the trichome base below the spike-like structure (Figure 3A-F). Occasionally, extensive hyphal growth was observed inside a broken trichome (Figure 3D). Growth on the palea surface without close interaction with trichomes (Supplementary Figure S3) and on/inside stomata was also common (Figure 3D+F).

**Figure 3.**
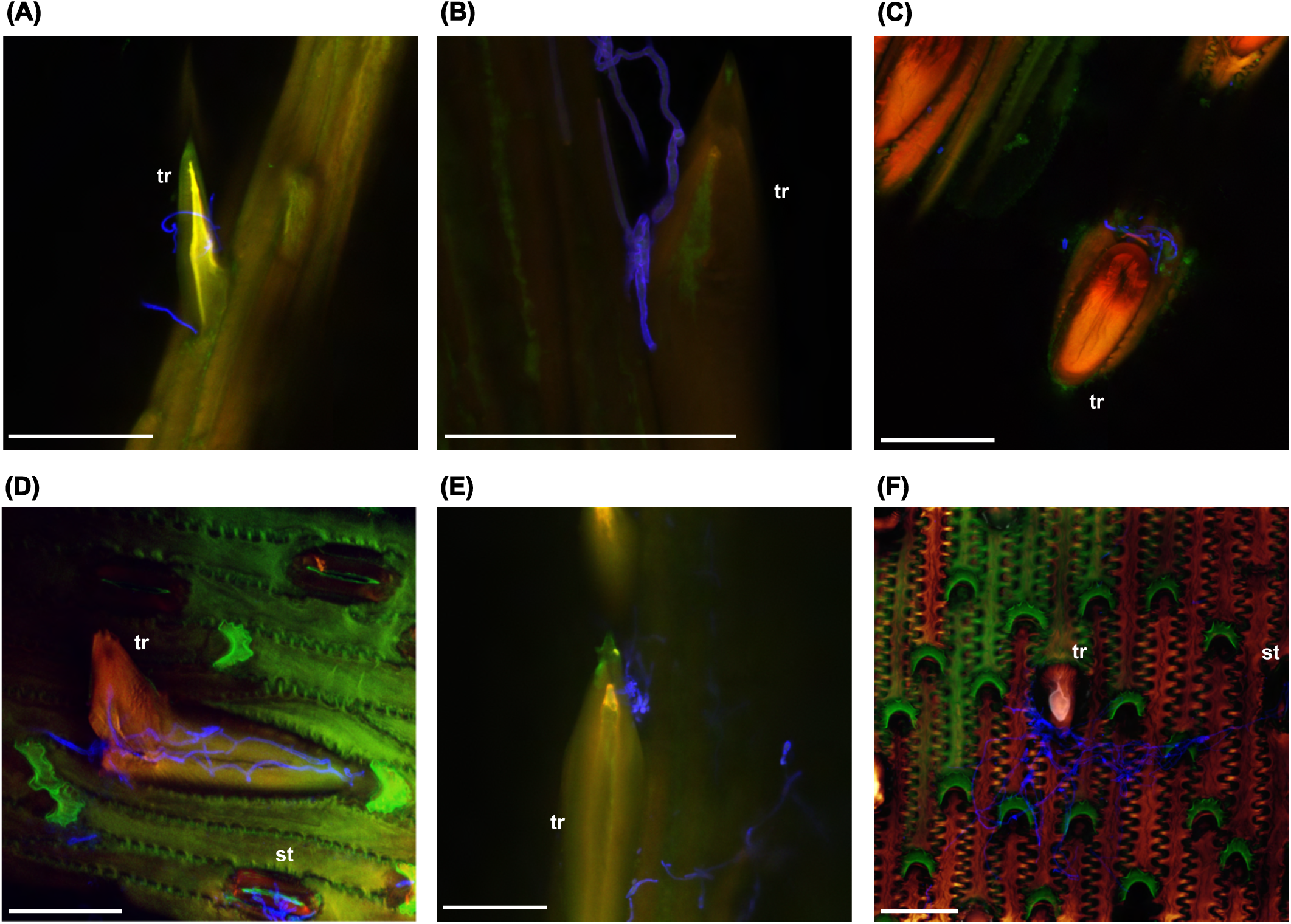
*F. graminearum* growth on trichomes (tr) and stomata (st) of oat paleas. **A.** 24 hpi. **B-D.** 48 hpi. **E-F.** 96 hpi. Fungal hyphae were stained using WGA-AF488 and are shown in blue. Maximum projections of CLSM overlay images. hpi: hours post inoculation. Scale bar: 50 µm.

## 4 Discussion

### 4.1 Environmental effects on FHB

In our study, the severity of *Fusarium* infection varied significantly depending on the fungal species and environment. Notably, most of the observed variance was explained by the environment, reflecting a strong influence of environmental conditions on *Fusarium* infection dynamics. These results align with other observations showing that *Fusarium* infection and mycotoxin contamination are strongly affected by temperature and moisture-related weather variables such as rainfall or relative air humidity especially during and after flowering (Hooker et al., 2002; Kriss et al., 2012; Parry et al., 1995; Xu et al., 2008). The production and distribution of conidia and ascospores from the spawn inoculum is also known to be strongly influenced by wind, temperature, rainfall and humidity, hence environmental conditions prior to infection are also important (Champeil et al., 2004; Crane and Bergstrom, 2014). In 2021, FP had the highest overall biomass, followed by FS, while FG had the lowest biomass. Environmental conditions may have favoured the other *Fusarium* species over FG. Furthermore, site-specific analysis showed that FG was highest in WL, which was in a region with higher rainfall than the other sites (Supplementary Table S9). The 2022 growing season was characterized by exceptionally dry conditions (ufz drought monitor, https://www.ufz.de/index.php?de=47252), which was accompanied by almost no infection by FS and FG. Only FP still showed considerable amounts of fungal biomass, suggesting that FP is more adapted to dry conditions than the other *Fusarium* species. This is consistent with the different environmental preferences reported previously for these *Fusarium* species. FP generally shows higher incidence after dry and warm weather conditions, whereas FG thrives in wetter and warmer environments (Doohan et al., 2003; Henriksen, 1999; Kriss et al., 2012; Meyer et al., 2021; Parikka et al., 2012; Turner et al., 1997; Xu et al., 2008). Furthermore, even within one field, considerable variability in *Fusarium* disease severity has often been observed, suggesting that very specific microclimatic conditions also play an important role (Oerke et al., 2010; Xu et al., 2008). In our study, weather data were not directly recorded at the field sites, but data from nearby regional stations were used to get a rough estimate of the environmental conditions. To gain a more detailed understanding of the specific preferences of the different *Fusarium* species in oats, measurements directly derived from the field that would better characterize the local microenvironment should be taken. In addition to environmental conditions, other factors such as tillage, previous crop and soil type can also affect the incidence of FHB, but to a lesser extent (Schaafsma and Hooker, 2007). In our study, these factors were the same for the three *Fusarium* species, but differed between the experimental sites and could also partly explain some of the observed site-specific differences. Taken together, our observations underline the necessity of evaluating *Fusarium* resistance across a large number of environments and replicates to ensure robust results.

### 4.2 *Fusarium* species-specificity and cross-resistance

FHB is caused by various *Fusarium* species. A generally effective, species-unspecific resistance would greatly help in breeding oat varieties with a reliably low risk of contamination with *Fusarium* mycotoxins. In our study, FP showed a significant positive correlation with FS biomass across host genotypes, whereas FG biomass was not correlated with either FS or FP. The positive correlation between FP and FS infection suggests partial cross-resistance, while the lack of correlation of FS and FP with FG rather suggests species-specific resistance mechanisms. In wheat, also a positive association between FP and FS was found and it was hypothesised that this could be due to the preferred production of microconidia in both species, and their relatively low competitiveness and aggressiveness (Oerke et al., 2010). In oats, mostly species-specific resistances and some general resistances have been reported. For example, Edwards (2009) found no correlation between HT-2 and DON levels in UK oat grain and therefore speculated that sources of resistance to one mycotoxin do not confer increased resistance to other mycotoxins. Similarly, no correlation of FHB resistance between FG- and FP-infested oat kernels was observed in a Canadian study (Tekauz et al., 2004). In a study by Hofgaard et al. (2022), the overall ranking of oat varieties based on HT-2/T-2 was not correlated with the ranking based on DON, although many varieties showed a similar ranking for both mycotoxins, suggesting some general resistance in addition to species-specific resistance.

Comparably, Herrmann et al. (2020) observed significant differences for DON and HT-2/T-2 levels, but also found a correlation between the ranking for both mycotoxins in some cases. Interestingly, the same oat UDP-glycosyltransferases are able to detoxify and confer resistance towards DON, NIV and HT-2 by heterologous expression in yeast, which would suggest that they could provide general resistance to different *Fusarium* species (Khairullina et al., 2022). However, the role of the different mycotoxins as virulence factors and the respective resistance mechanisms are still largely unknown in oats. A possible reason for species-specificity of resistance could be the different epidemiology of the *Fusarium* species. In addition to distinct preferences for environmental conditions, the timing of the influence of weather conditions on infection may also differ between *Fusarium* species (Hjelkrem et al., 2017, 2018). Some morphological or phenological characteristics, such as plant height, earliness or anther retention, may also explain some of the differences in resistance, but were not or only partly assessed in our study.

Heritability was markedly low for FP, where no significant genotype effect was found. This indicates a lack of significant genetic variability or extreme dependence on environmental conditions particularly for this *Fusarium* species. Another study also found no significant differences in FP DNA levels between oat genotypes but rather location-specific effects on FP abundance (Martin et al., 2018). The reason for the difference in genotypic effect sizes between the *Fusarium* species is unclear, but could be related to species-specific resistance mechanisms as well. In our study, oat genotypes had small differences in plant height and earliness, which are factors known to influence *Fusarium* resistance ranking. The possibility that greater differences in these factors would result in significant differences in resistance to FP should be investigated. The lack of genetic effect for resistance to FP is concerning as this makes breeding for FP resistance particularly difficult. This is especially problematic because, although FP is considered a rather weak pathogen, its prevalence has been increasing and it is often found to be one of the most abundant *Fusarium* species in oats (Karlsson et al., 2022; Kuchynková and Kalinová, 2021; Martin et al., 2018; Parikka et al., 2012; Parry et al., 1995; Stenglein, 2009; Valverde-Bogantes et al., 2020; Xue et al., 2019). Also in our study, FP biomass was overall much higher than FG and FS biomass.

Although some correlation could be found for resistance to different *Fusarium* species, it is questionable whether there is sufficiently strong cross-resistance in oats. Therefore, separate tests are most likely required to reliably determine resistance to different *Fusarium* species. This makes breeding for durable *Fusarium* resistance even more challenging, as the occurrence of *Fusarium* species is expected to constantly shift and previously marginal species may become more important due to climate change (Parikka et al., 2012; Hietaniemi et al., 2016; Moretti et al., 2019; Valverde-Bogantes et al., 2020; Karlsson et al., 2022). Nevertheless, some genotypes in our study showed consistent resistance across *Fusarium* species and could be useful candidates for breeding programmes. Their resistance phenotypes should however be validated in additional environments due to the high variability. Whether their resistance is based on broadly effective mechanisms or on a combination of different resistance sources for each *Fusarium* species is not clear and should be investigated.

### 4.3 Variation of trichome size and density

This study revealed a significant variation in hull trichome density and size among oat genotypes, and the genotypic differences were stable across environments. This is reflected by a very high heritability, suggesting that this trait is strongly influenced by genetic factors.

Furthermore, the trichome traits were highly correlated between lemma and palea, suggesting a coordinated regulation of trichome development across these organs. This also means that the laborious trichome phenotyping process could be simplified by focusing on only one of these organs and using less environments in future studies.

The observation that only prickle trichomes were present on the lemma and palea is consistent with findings in wheat, where prickle trichomes were also the only type on the florets (Sun et al., 2024). In contrast, florets of two-row barley display distinct, dome-shaped trichomes, while prickle-like trichomes are present mostly on six-row barley (Imboden et al., 2018).

In wheat and rice, several loci that control trichomes on florets and/or leaves have been mapped (Angeles-Shim et al., 2012; Chen et al., 2021; Wu et al., 2021; Luo et al., 2022; Fan et al., 2023). In oats however, the genetic basis of hull trichome formation has not been investigated until now. It would be interesting to use contrasting genotypes identified in this study to develop a segregating population to map genetic loci for trichome development in oats as well.

### 4.4 Trichomes in *Fusarium*-host interactions

We found a significant positive correlation between FG biomass and trichome density, meaning that oat genotypes with fewer trichomes on the hulls were generally more resistant, suggesting that a higher number of trichomes may facilitate colonization by FG. This is consistent with results from wild *Avena* species, where trichome density and *Fusarium* infection were positively correlated as well (Gagkaeva et al., 2017). Interestingly, in wheat, genetic loci controlling trichome length and density overlap with QTLs for FHB resistance, which also points towards a close relationship between trichomes and *Fusarium* infection (Häberle et al., 2009; Zheng et al., 2021). In contrast, Duba et al. (2019) found that leaf trichome density was higher in resistant than susceptible wheat lines, rather suggesting a protective role of trichomes. It should however be noted that the observed correlation with FG is based on infection data from a relatively small number of environments and should therefore be interpreted with caution. Nevertheless, the involvement of trichomes in FG infection in oats is further supported by our microscopic observations, where FG hyphae frequently accumulated in close contact with trichomes. FG hyphae were also growing on stomata, indicating multiple colonization routes. This is in line with findings from other *Fusarium*-host systems. In maize, FG hyphae penetrate through the tip of prickle trichomes (Nguyen et al., 2016), while in wheat, the base is penetrated (Sun et al., 2024). In barley, also the base of prickle trichomes is invaded, while trichomes remain intact (Imboden et al., 2018; Linkmeyer et al., 2013). In wheat, invasion is more aggressive and trichomes are destroyed by secretion of cell wall-degrading enzymes during the infection process (Sun et al., 2024). Furthermore, entry through stomata is a common infection route for *Fusarium* in various hosts as well (Beccari et al., 2011; Kang et al., 2000; Linkmeyer et al., 2013; Pritsch et al., 2000). We also observed growth of FG hyphae on broken trichomes, which may indicate that the fungus is actively destroying trichomes for penetration, similar to what happens in wheat. A more detailed study of the microscopic interaction of hyphae and trichomes as infection progresses is required to clarify this and to determine whether the hyphae specifically penetrate the trichome base in oats as well.

There are several possible reasons for the important role of trichomes for *Fusarium* infection. Firstly, trichomes could serve as attachment points for spores, preventing them from being easily carried away by wind or rain. Alternatively, trichomes could provide a microclimate favourable to fungal growth, such as increased humidity due to water droplets adhering to them (Calo et al., 2006; Imboden et al., 2018). In grasses, trichomes accumulate silica, similar to other cell types that are also preferred entry sites for *Fusarium* hyphae such as vascular bundles, stomata and silica cells. This suggests that the fungus may be specifically attracted to high silica levels (Rittenour and Harris, 2010; Boenisch and Schäfer, 2011; Imboden et al., 2018). The trichome base also appears to be a particularly vulnerable site for *Fusarium* infection (Sun et al., 2024). Differences in cell wall composition between trichomes and other epidermal cells may therefore provide another explanation. In contrast to FG, no significant correlation was identified between trichomes and FS or FP biomass in our study. This may be due to the lower heritabilities of FS and FP infection severity, but may also indicate species-specific interactions with trichomes in oats. Depending on the actual function of trichomes in the infection process, different explanations are possible. *Fusarium* species have different climatic preferences, so the microclimate provided by the trichomes may only be conducive to the growth of FG, but not to the other *Fusarium* species tested. Furthermore, the conidia of FG, FS and FP differ considerably in size and shape. The large lunate-shaped macroconidia of FG might be more efficiently trapped by the prickle trichomes, whereas the smaller and more rounded micro- and mesoconidia of FP and FS are not as efficiently attached (Palicova et al., 2024). It is also conceivable that not all *Fusarium* species possess the enzymes necessary to degrade the oat trichome cell walls.

However, in contrast to our results, in barley, penetration of trichomes was found to be an important infection route for FS as well (Linkmeyer, 2012), and to our knowledge, no *Fusarium* species-specificity for the role trichomes has been described previously. Whether there are species-specific interactions with trichomes in oats or whether the lack of correlation is rather due to the low heritability in the field trials needs to be clarified in future studies. For this, infection results from controlled experiments will be helpful, as the involvement of trichomes may be easier to assess in the absence of highly variable environmental conditions.

In conclusion, it might appear promising that by the use of oat genotypes with fewer trichomes, FG infection could be reduced. However, it should be taken into account that trichomes also serve important positive functions such as reducing insect feeding and increasing resistance to UV radiation (Mauricio and Rausher, 1997; Peeters, 2002; Yan et al., 2012; Zhang et al., 2021).

Consequently, the reduction of trichome density might not be an advisable breeding target. As the trichome base seems to be the most vulnerable part, Sun et al. (2024) suggest that breeding efforts should be directed towards strengthening the trichome base to make penetration more difficult.

Another potential approach might be the targeted expression of antifungal compounds in trichomes to inhibit fungal infection. In *Arabidopsis*, resistance to *Botrytis cinerea* could be increased by trichome-specific expression of a *Trichoderma* α-1,3-glucanase, an enzyme that hydrolyses the cell wall of various fungi (Calo et al., 2006). However, how useful these approaches would be in practical applications is uncertain and remains to be explored.

## 6 Author Contributions

SS: Formal analysis, Investigating, Visualization, Methodology, Writing – original draft, Writing – review & editing. CR: Investigating, Methodology, Writing – review & editing. KO: Resources, Writing – review & editing. StB: Resources, Writing – review & editing. SoB: Resources, Writing – review & editing. AvT: Resources, Writing – review & editing. MH: Conceptualization, Funding acquisition, Formal analysis, Resources, Writing – review & editing.

## Acknowledgments

We would like to acknowledge the imaging facility of the Julius Kühn Institute for Epidemiology and Pathogen Diagnostics for use of the Leica TCS SP8 confocal laser-scanning microscope and are grateful to Yvonne Becker for helpful support on sample preparation and image acquisition. DeepL Write (beta) was used to support language editing.

## Supplementary

**Supplementary Figure 1.**
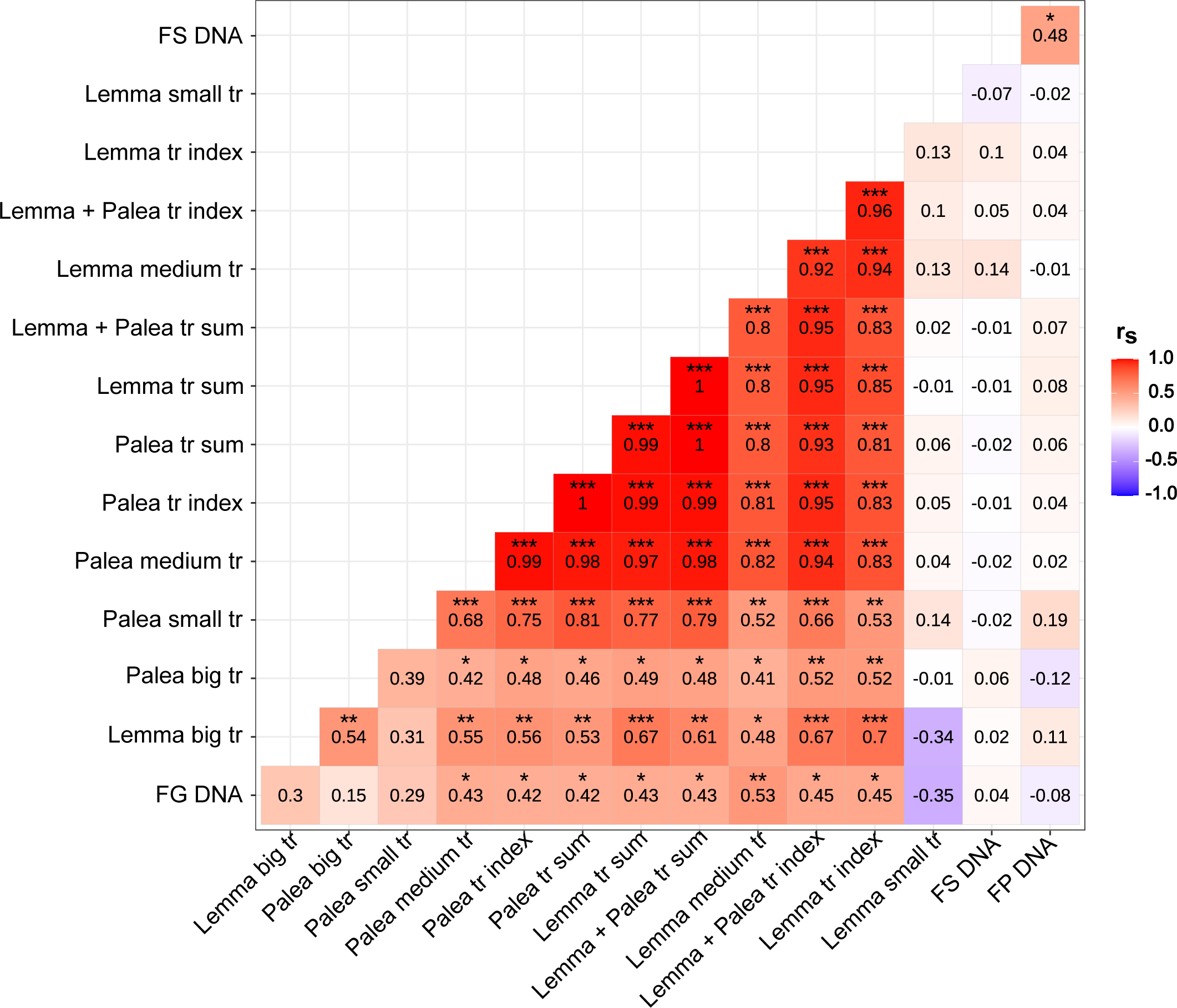
Correlation matrix of trichome counts (sum) and trichome indices and *Fusarium* biomass. Fungal biomass was assessed via quantification of fungal DNA per kg oat seeds. BLUEs were used for calculation of Spearman rank correlation coefficients. Asteriscs indicate p value significance levels with p < 0.001: ***; p < 0.01: **; p < 0.05: *. tr: trichome; FG: *F. graminearum*; FP: *F. poae*; FS: *F. sporotrichioides;* rs: Spearman - rank correlation coefficient.

**Supplementary Figure 2.**
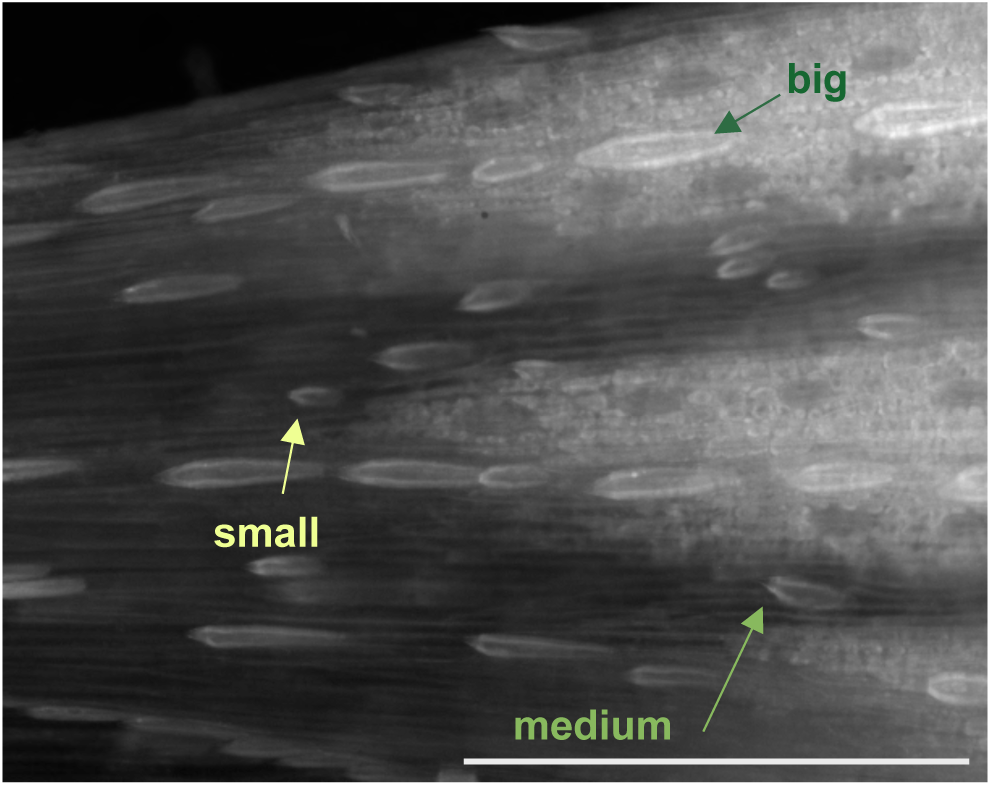
Examples of trichomes for each size class. Scale bar: 500 µm.

**Supplementary Figure 3.**
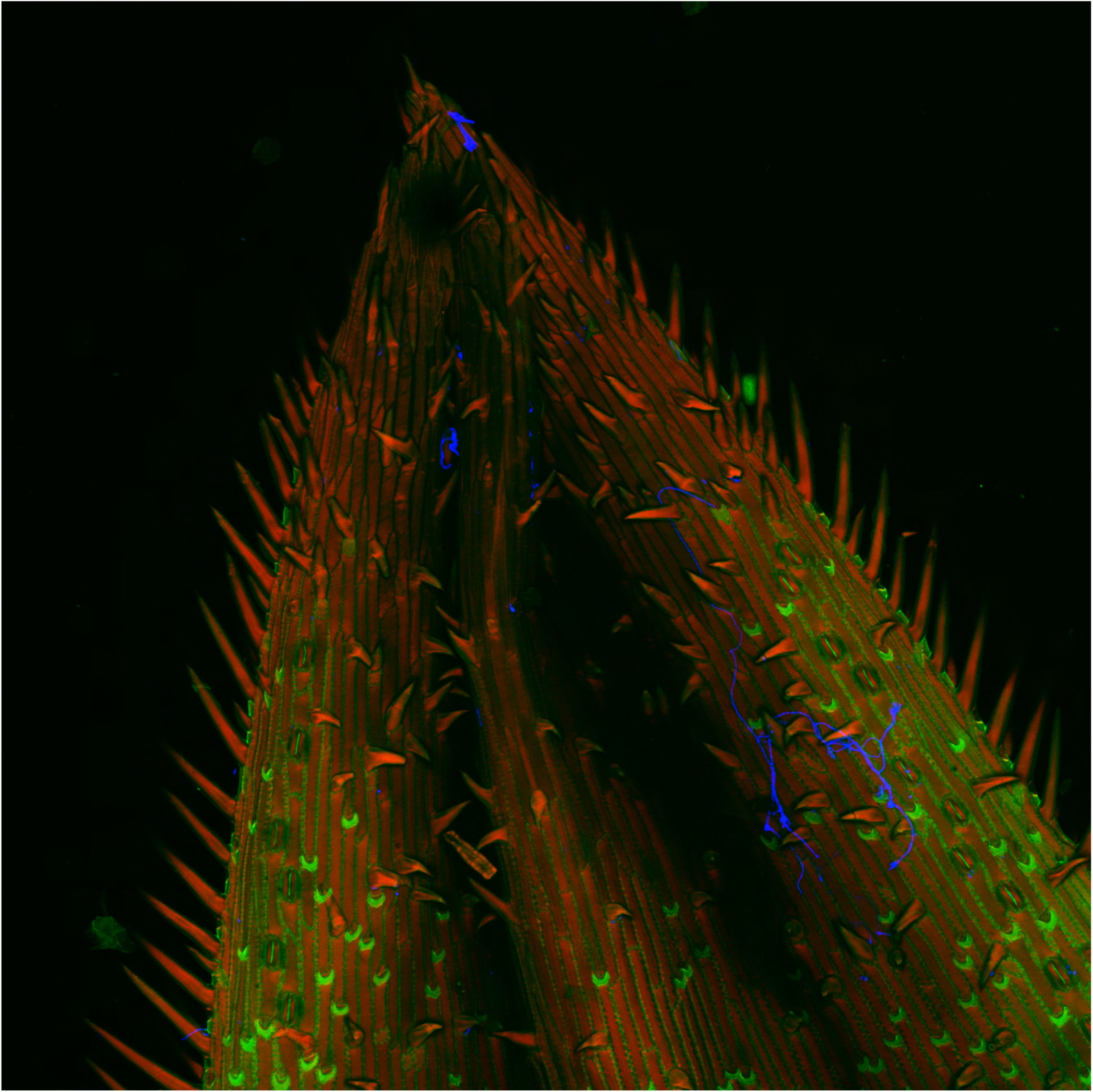
Overview image of *F. graminearum* growth on oat palea 96 hpi. Fungal hyphae were stained using WGA-AF488 and are shown in blue. Maximum projection of CLSM overlay images. hpi: hours post inoculation.

